# Regional alignment with sex-typical brain architecture, maternal health and environmental enrichment predict psychiatric symptoms in early life

**DOI:** 10.64898/2026.09.27.754771

**Authors:** Raluca Petrican, Leonardo de Pascalis, Christopher Mugatroyd, Victoria Fallon, Sophie Pendered, Tamara Vaz

**Affiliations:** Institute of Population Health, Department of Psychology, University of Liverpool, Bedford Street South, Liverpool, L69 7ZA, UK; Department of Psychology, University of Bologna, Bologna, Italy; Department of Life Sciences, Manchester Metropolitan University, Manchester, UK; Child and Adolescent Health Psychology, Department of Psychology, University of Zurich

**Keywords:** Functional connectivity, sex differences, perinatal adversity, cognitive enrichment, autism, ADHD, behavioral disorders

## Abstract

Robust sex differences in brain morphology and function exist from birth onwards. While traditionally conceptualized as categorical, they were recently shown to be best described along a female-to-male continuum. Here, we probed the under-explored implications of this proposal for understanding early life vulnerability to psychiatric conditions with sex-biased prevalence. We analyzed environmental, clinical and resting state functional neuroimaging data from 411 participants in Developing Human Connectome Project (dHCP) and 214 children from the Oregon ADHD 1000 study. In the dHCP, we identified complementary patterns of neonatal alignment with sex-typical young adult architecture, which overlapped areas rich in nicotinic cholinergic receptors and predicted the same psychiatric symptoms across sexes in toddlerhood. Specifically, for association (compared to sensorimotor) areas, greater alignment with female-typical functional connectivity (FC) patterns foreshadowed higher anxiety, whereas greater alignment with male-typical FC patterns forecasted autism-related problems. The latter effect was weaker among toddlers with an older epigenetic age at birth and exposure to fewer maternal health problems, but greater environmental enrichment. In the independent child sample, patterns of alignment with sex-typical FC, overlapping those observed in neonates, distinguished a history of depression and generalized anxiety (greater alignment with female-typical FC) from a history of separation anxiety, behavioral problems and current ADHD (greater alignment with male-typical FC). Our study speaks to the psychiatric relevance of sex-related functional brain architecture from birth onwards, its value in identifying vulnerable youths and its promise in monitoring the success of environmental interventions.

---

Sex differences in brain morphology and function are present from birth and, with advancing age, they become increasingly salient and predictive of distinguishable mental health trajectories (Bangasser & Cuarenta, 2021; Dhamala et al., 2023; Eme, 2016; Gilmore et al., 2018; Solmi et al., 2022). Neuroscience research has traditionally treated sex as dichotomous, a conceptualization challenged by recent findings showing that the human brain is best described along a female-to-male differentiation continuum (Zhang et al., 2021), with each individual brain containing some features that tend to be observed most often in males, some that are more common in females, and others that are equally distributed across sexes (Joel et al., 2015). Thus, taking into consideration each brain’s unique tapestry of male and females features could be key to personalizing psychiatric detection and intervention, particularly for disorders showing sex-biased prevalence (Bangasser & Cuarenta, 2021; Dhamala et al., 2023; Eme, 2016; Hampel et al., 2023).

To date, investigations of sexual differentiation in brain morphology and function tended to sample individuals from late childhood onwards. Consequently, very few could disentangle the impact of sex from that of gender, since such distinction requires thorough assessment of both (e.g., Dhamala et al., 2024). Here, we took a step towards addressing this issue by probing the psychiatric relevance of neural functional coupling (FC) patterns connected to biological sex in a sample of newborns (N = 411) from the Developing Human Connectome project (dHCP) who had yet to be exposed to any gendered experiences. Our first objective was to elucidate whether regional patterns of neonatal alignment with sex-typical FC patterns would foreshadow, over the first two years of life, symptomatology of disorders with a sex-biased prevalence (i.e., depression, anxiety vs autism spectrum disorder [ASD], attention deficit hyperactivity disorder [ADHD], oppositional defiant disorder [ODD]) (Bangasser & Cuarenta, 2021; Dhamala et al., 2023; Eme, 2016; Loomes et al., 2017). We were primarily interested in whether, across sexes, neonatal alignment with female-typical FC patterns would predict internalizing symptoms, whereas alignment with male-typical FC patterns would forecast externalizing and/or ASD-relevant problems.

Our second objective was to examine the malleability of the relationship between neonatal alignment with sex-typical FC patterns and psychopathology at 18 months by probing its susceptibility to the impact of variables implicated in neurodevelopment and psychological functioning during the first two years of life. We thus tested the influence of infant-related characteristics (i.e., gestational age [GA] at birth, neonatal epigenetic age) and of perinatal environmental factors related to the mother (i.e., physical and mental health) and to the home environment (i.e., cognitive enrichment, that is, the child’s exposure to mentally stimulating toys, books, and activities in the home, and parental behaviours that encourage engagement in mentally stimulating activities and learning of new skills (Einziger et al., 2021; Einziger et al., 2019)). Premature birth and thus, younger GA, has been robustly linked to delayed or atypical neurocognitive development (Ball et al., 2013; Brandstaetter et al., 2025; Cheong et al., 2017; McKinnon et al., 2023; Menu et al., 2025). Relatedly, epigenetic aging, measured via differential DNA methylation (DNAm) patterns, varies between psychiatric disorders with sex-biased prevalence (e.g., accelerated aging in internalizing disorders (Kuan et al., 2021; Protsenko et al., 2021)) and mediates prenatal environmental effects (e.g., maternal mood, socioeconomic status) on the offspring’s health (Abrishamcar et al., 2024; Choudhary et al., 2024), including neuropsychiatric vulnerability (Kallak et al., 2022; Kotsakis Ruehlmann et al., 2023). As for the perinatal environment, a burgeoning cross-species literature attests to the critical contribution of cognitively stimulating environments (i.e., perceptually rich and novel environments that foster learning) to reversing the sequalae of perinatal adversity (e.g. poor maternal health, Wang et al., 2025) or caregiving (You et al., 2025) on neurogenesis and functional brain network maturation.

Conversely, an extensive body of work testifies to the key role of maternal health in their offspring’s development. For instance, maternal medical problems, particularly metabolic (e.g., diabetes), cardiovascular (e.g., hypertension) and immune (e.g., infections) disorders predict deviations from typical brain maturation (Katsuki et al., 2021; Niskanen et al., 2025), heightened risk for neurodevelopmental conditions (Duan et al., 2025; Rodolaki et al., 2023), and sex-dependent vulnerability to internalizing (females) vs externalizing (males) symptoms in childhood (Lipner et al., 2024). Likewise, maternal depression, anxiety and adversity exposure (inferred from hormonal indices) during gestation and toddlerhood show robust connections with atypical brain development (Gilmore et al., 2018; Mareckova et al., 2020), including sexually dimorphic effects (Kumpulainen et al., 2023; Sandman et al., 2018), particularly on the limbic system (Manning et al., 2025), as well as poorer sociocognitive development from birth to adolescence (Girchenko et al., 2025).

Our third objective was to probe whether the identified patterns of neonatal alignment with sex-typical brain architecture (from Objective 1) align with existing functional hierarchies of brain maturation (Margulies et al., 2016), specifically, the sensorimotor-association (S-A) axis along which typical neurodevelopment and associated sex differences reportedly unfold (Luo et al., 2024; Petrican et al., 2025a; Shanmugan et al., 2022; Sydnor et al., 2025; Sydnor et al., 2021; Sydnor et al., 2023) and the visual-sensorimotor axis, which constitutes the main functional axis at birth (Gilmore et al., 2018; Lariviere et al., 2020). To delineate potential pharmaceutical targets and situate our findings in relation to existing neuropsychiatric interventions, we also explored the neurochemical correlates of our neonatal brain patterns. We focused on the receptor and transporter density maps associated with the major neurotransmitter systems described in Hansen et al.(2022), which are critically implicated in brain and cognitive maturation, as well as vulnerability to neurodevelopmental disorders and psychopathology more broadly (i.e., acetylcholine [Ach], (Ananth et al., 2023; Ballinger et al., 2016), dopamine [DA], (Li et al., 2025), GABA/glutamate [GLU], (Basu et al., 2023; Markicevic et al., 2020), norepinephrine [NE], (Bang et al., 2023; Bari et al., 2020; Dahl et al., 2022; Mather et al., 2016), serotonin [5-HT], (da Cunha-Bang & Knudsen, 2021; Erritzoe et al., 2023; Svensson et al., 2021).

Our final objective was to investigate whether overlapping patterns of alignment with female-vs male-typical FC could differentiate between internalizing and externalizing symptoms both in toddlerhood and in later childhood. To this end, we probed the spatial overlap between the neonatal brain patterns identified in the dHCP sample (see Objective 1) and the brain maps tracking alignment with sex-typical FC as a function of internalizing (depression, anxiety) vs externalizing (ADHD, ODD) symptoms in a community child sample.

## 2. Methods

### 2.1 Cohorts

Our primary analyses featured neonatal and 18-month follow up data from 411 participants in the dHCP study, of whom 313 were born to term (152 females) and 98 were born preterm (46 females). Our secondary analyses focused on baseline data from 214 youths (aged 7-12 years, 85 females) from the Oregon ADHD 1000 study, a longitudinal ADHD case-control community sample (Nigg et al., 2023), which showed a relatively substantial lifetime history of internalizing and externalizing disorders (see Table 1). Biological sex assignments, as reported by the parents/caregivers, were corroborated via single nucleotide polymorphism (SNP) analyses. At the time of testing none of the participants were on any medication. Detailed description of both samples, including genetic sex assignment, is included in the Supplemental Information (SI). Figure S1 contains the selection criteria for the dHCP and Oregon samples.

### 2.2. Measures

The measures and the characterization of the sex-related FC gradient, which are briefly described below, are further detailed in the SI.

#### 2.2.1 Health Assessments

##### dHCP (18-month follow-up)

The toddlers’ psychological health was measured via parental responses on the Child Behavior Checklist (CBCL) for ages 1.5-5 (Achenbach & Ruffle, 2000) which yields scores on five DSM-oriented scales for Affective Problems, Anxiety, Autism Spectrum Disorder (ASD)/Pervasive Developmental Problems, Attention Deficit Hyperactivity Disorder (ADHD) and Oppositional Defiant Disorder (ODD). Maternal depression and anxiety were derived from self-ratings on the 7-item depression and the 3-item anxiety subscales from the Edinburgh Postnatal Depression Scale (Cox et al., 1987), which can adequately distinguish between maternal risk for major depression and anxiety disorders (Phillips et al., 2009). Self-reported maternal medical history covered serious conditions that preceded (e.g., malignancy, diabetes, asthma) or occurred during the pregnancy (e.g., pre-eclampsia, gestational diabetes, fever during the delivery).

##### Oregon sample

At baseline (ages 7-12 years), current diagnosis of ADHD and lifetime diagnosis of MDD, anxiety disorders (separation, social, generalized) and ODD were established based on parental responses in a semi-structured clinical interview with a trained Master’s-degree level clinician (Kiddie Schedule for Affective Disorders and Schizophrenia [KSADS](Kaufman et al., 1997)). A clinical diagnosis of ASD was not present and conduct disorder cases (N =3) were not sufficiently represented in the sample with complete data on our variables of interest.

#### 2.2.2 Evaluation of the home environment

Availability of cognitively enriching experiences (e.g., museum visits) and toys (e.g., toys that stimulate number knowledge) was evaluated via parental responses on a version of the Cognitively Stimulating Parenting Scale (Wolke et al., 2013). The dHCP study team had revised the scale to include toddler access to technology (e.g., iPhones and Apps) (Bonthrone et al., 2021).

#### 2.2.3 Epigenetic aging

Epigenetic age was estimated from saliva extracted DNA using the Pediatric-Buccal-Epigenetic (PedBE) clock introduced by McEwen et al. (2020) and validated for ages 0-20 years. Similar to McEwen et al. (2020), after adjusting for sex and postnatal age (in days) at scan, we found a positive partial correlation between the PedBE age estimate and gestational age at birth (in days), Spearman’s *rho* =.22, permutation-based *p* = 2 x 10^-5^.

### 2.3 Functional Neuroimaging Data: Sex-Typical FC Strength Gradient

Alignment with sex-typical FC strength patterns was estimated from 15 minutes of task-free data collected during sleep (dHCP) or awake rest (Oregon). Pairwise FC indices were computed based on two functional atlases (one featured in the main analyses (Schaefer et al., 2018), the second included for replication (Gordon et al. 2016) which had been developed in adults, but used with both neonates (Tooley et al., 2024; Yin et al., 2025) and children/adolescents (Hagler et al., 2019; Petrican et al., 2024; Petrican et al., 2025a). This approach facilitated comparisons with prior work conducted across the life course (see SI for details on infant-specific atlas alignment). For each dHCP neonate or Oregon youth, similarity to sex-typical FC was estimated as correlation-based alignment with the FC strength patterns observed among the female and male participants (aged 22-30 years) in the Human Connectome Project (HCP) (see Petrican et al., 2025a and SI for details). Pubertal hormones play a reportedly critical role in the development of brain regions that are sexually dimorphic across species and implicated in the pathophysiology of disorders with sex-asymmetric prevalence (Ahmed et al., 2008; Piekarski et al., 2023; Schulz & Sisk, 2016). Therefore, because the HCP participants had reached reproductive maturation, we expected them to show maximal differentiation in sex-typical FC patterns.

To identify sex-typical FC strength patterns in the HCP, we regressed each of the 44850 parcel-to-parcel FC indices against sex and a number of confounds which had been linked to individual differences in FC and on which complete matching between sexes is not feasible, specifically, age, income, employment status, race, handedness, average in-scanner motion, intelligence and psychopathology. For each parcel-to-parcel FC index, the *t*-statistic corresponding to “sex” was taken as a measure of the robustness with which the respective FC index differentiates between men and women. We focused on the t-statistic, rather than the beta weight corresponding to “sex”, in order to safeguard against prioritising FC indices that show large sex differences in a subsample, but not the entire HCP group (i.e., FC indices with high absolute value beta weights, but also high standard errors, suggestive of greater between-individual variability and, thus, lower robustness in distinguishing between sexes). Nonetheless, we verified that the *t*-statistic and corresponding beta weight value associated with “sex” for a given parcel-to-parcel FC index were very strongly correlated (median *r* of.96, range:.79-.995), as expected given the intentional selection of a relatively homogenous group of HCP participants to act as reference for the sex-related FC gradient analyses.

#### 2.3.1. Alignment with the HCP sex-related FC gradient in the dHCP and Oregon samples

For each dHCP or Oregon participant, relative alignment with female-vs male-typical FC strength patterns was computed as the rank-order correlation coefficient between each column of their own FC matrix (i.e., each parcel’s FC with each of the remaining parcels in the atlas) (Figure 1-H) and the corresponding column in the HCP *t*-statistic matrix (Figure 1-G). A positive correlation coefficient indicates that the regional FC strength profile from a dHCP or Oregon participant aligns with the ascending female-to-male FC strength gradient from the HCP (implying a more masculine FC strength spatial profile). A negative correlation coefficient indicates that the regional FC strength profile from a dHCP or Oregon participant aligns with the ascending male-to-female FC strength gradient from the HCP (implying a more feminine FC strength spatial profile). Because the HCP *t*-statistic matrix is based only on young adult FC profiles, anchored at one end in female prototypicality and, at the other, in male prototypicality, both positive and negative rank order correlation coefficients indicate similarity to a young adult brain profile.

**Figure 1.**
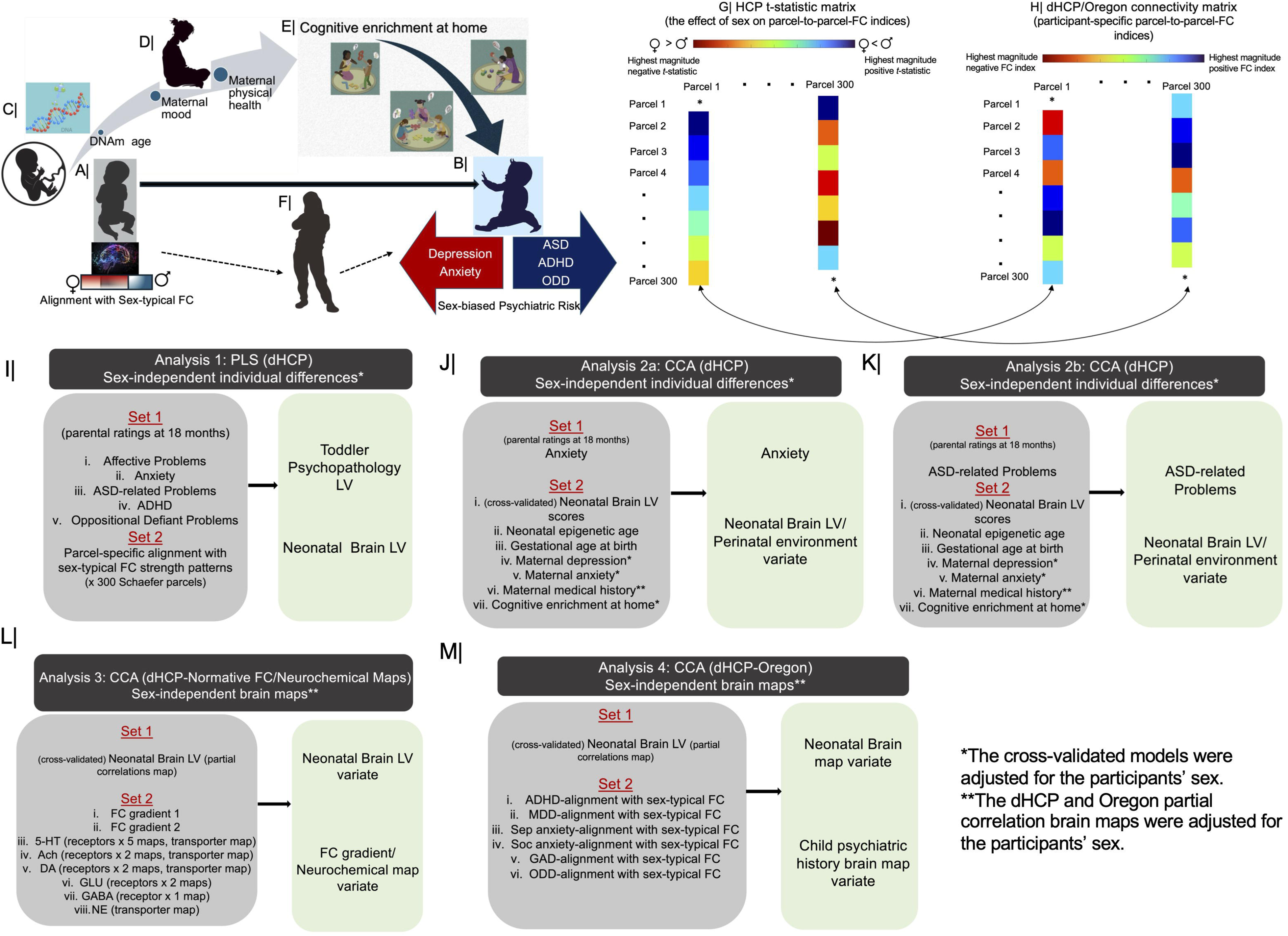
Overview of our research objectives and analytic strategy. Top panels. Here, we probed the psychiatric relevance of neonatal and child alignment with sex-typical brain architecture. Specifically, we tested whether individual differences in neonatal alignment with female-vs male-typical FC (panel A) predict, in toddlerhood, vulnerability to psychiatric disorders that show sex-biased prevalence (i.e., depression, anxiety vs ASD, ADHD, ODD) (panel B). We further investigated whether the predictive power of alignment with sex-typical brain architecture is impacted by perinatal factors related to the infant (i.e., DNAm age) (panel C), the mother (i.e., physical health up to and including the pregnancy, depression and anxiety 18 months after the birth) (panel D) and/or the home environment (i.e., availability of cognitively stimulating toys, activities) (panel E). Finally, we examined the developmental stability of the identified neonatal patterns (from panel A), specifically, their predictive power for similar psychiatric symptoms in toddlerhood and childhood (panel F). In panel (F), the arrows indicate the hypothesized directionality of the effects, specifically, that some of the regional patterns of alignment with sex-typical FC, which are present from birth (panel A), correlate with the same mental health symptoms in toddlerhood (panel B) and childhood (panel F). The arrows are in dashed line because the cross-sectional analyses of the Oregon data did not allow any tests of the hypothesized temporal sequence. Panels G-H depict the analytic steps through which alignment with female-vs male-typical FC patterns was estimated. *T*-statistic values corresponding to self-connections (panel G) or FC indices corresponding to self-connections (panel H) were set to zero. Bottom panels. Analysis 1: PLS related regional patterns of neonatal alignment with sex-typical FC to parental ratings of affective problems, anxiety, ASD-related challenges, ADHD and ODD. Analysis 2a/b: CCA linked individual differences in the cross-validated neonatal brain LV from Analysis 1, epigenetic age, gestational age at birth, maternal health (medical history, depression, anxiety) and cognitive enrichment at home to individual differences in the two psychiatric functioning scales robustly correlated with sex-related brain architecture in Analysis 1 (anxiety, ASD). Analysis 3: CCA related the cross-validated neonatal brain LV map from Analysis 1 to the FC gradient maps (gradient 1: unimodal-to-transmodal areas; gradient 2: visual-to-sensorimotor) (Margulies et al., 2016) imported from neuromaps (Markello et al., 2022), as well as the receptor and transporter density maps for major neurotransmitter systems, created by Hansen et al. (2022) and downloaded from https://github.com/netneurolab/hansen_receptors. Analysis 4: CCA probed the relationship between the cross-validated neonatal brain LV map from Analysis 1 and the brain maps tracking alignment with sex-typical FC as a function of a current diagnosis of ADHD, lifetime diagnosis of MDD, anxiety ([separation, social, generalized], ADHD and ODD in the Oregon sample. The Oregon brain maps were estimated with partial Spearman’s correlations, adjusted for the confounders listed under “Control variables for the dHCP and Oregon samples”. LV = latent variable. FC = functional connectivity. ADHD = attention deficit hyperactivity disorder. ASD = autism spectrum disorder. MDD = major depressive disorder. Sep = separation. GAD = generalized anxiety disorder. ODD = oppositional defiant disorder. 5-HT =serotonin. Ach = acetylcholine. DA= dopamine. GABA = gamma-aminobutyric acid; GLU = glutamate. NE = norepinephrine.

#### 2.3.2 Validation of the HCP sex-related FC gradient

The HCP sex-related FC strength gradients were validated in the dHCP and Oregon samples. Specifically, we confirmed that, in each of the two samples, male participants showed a more positive average correlation between their own FC matrix and the female-to-male FC strength gradient defined in the HCP sample relative to the female participants (cf. Figure 1-G, H). The validation and the results reported below were replicated with an alternate atlas and HCP sex-related FC gradient which further controlled for brain size and female menstrual cycle characteristics, variables that reportedly impact structural brain characteristics (Küchenhoff et al., 2024). In supplemental analyses (Figures S5-6, A-F), we confirmed the assumption underlying our comparisons with the sex-related FC gradient from the HCP sample, which is that alignment with sex-typical FC profiles emerges gradually from birth onwards, with young adults constituting the most appropriate reference group because they reached reproductive maturation and would thus show maximal differentiation in sex-related FC. Further support for this reasoning came from supplemental tests showing that, while female-to-male FC strength gradients using the dHCP or Oregon data as a reference perform well at differentiating between sexes within the respective age group (Figures S5-6, G, H), they cannot discriminate between sexes across age groups (permutation-based *p*-values >.27).

A step-by-step description of the analyses summarised above is included in the SI.

### 2.4 Statistical Analysis

We leveraged two data-driven multivariate analysis techniques, partial least squares (PLS) correlation analysis and canonical correlation analysis (CCA), which explain commonalities in two sets of variables (Set 1 and Set 2 in Figure 1, I-M) by creating linear combinations, dubbed “variates” (CCA) or “latent variables” (PLS), from the observed variables in each set (Hair et al., 2014; Krishnan et al., 2011; McIntosh & Misic, 2013; Mihalik et al., 2022). Both were implemented in Matlab (2025b) using a 10-fold cross-validation procedure. Statistical significance was assessed through permutation testing (100,000 samples), whereas bootstrapping (100,000 samples) was used to evaluate the reliability of within-set relationships (99% confidence intervals [CIs] for canonical loadings [CCA] and saliences [PLS]) and between-set relationships (95% CIs for cross-loadings [CCA], observed behavioral variables-brain latent variable [LV] correlations [PLS], correlations between the extracted variates [CCA] or LVs [PLS]). We conducted 5 multivariate analyses (Figure 1, I-M) for which we ran the technical replications reported in the SI. Statistical significance in the discovery and corresponding cross-validation (aka “simulated replication”) analyses was assessed only for the LV (PLS) or variate (CCA) pair explaining the highest amount of covariance between the two sets of observed variables in each of the 5 discovery analyses. There was no need to correct across analyses because no more than 3 multivariate models considered the same type of data.

### 2.5 Control variables for the dHCP and Oregon samples

To facilitate interpretability of the model estimates, we ran the discovery CCA and PLS analyses on the raw/non-residualized data. To demonstrate that our results are robust against variations in known confounders, the CCA and PLS cross-validation analyses were adjusted for the participants’ sex assigned at birth, chronological age, race, maternal education level (as a well-supported proxy for socioeconomic status (Cantarutti et al., 2017) which, unlike income, was available in both target samples), and summary in-scanner motion per participant (see SI for details). Additional controls applied only to the dHCP data are detailed in the corresponding Results sections below.

### 2.6 Design Considerations

The present study has a cross-sectional correlational design and includes a cross-temporal correlational component for the dHCP sample. For the two target developmental samples, we used the maximum number of participants with available data on all the variables of interest. The inter-related nature of our analyses precluded use of participants with partially available data. The dHCP and Oregon participant selection process is depicted in Figure S1. The size of our analyzed dHCP sample matches and, in some cases, exceeds the sample sizes used in similar investigations of brain development during the first two years of life (Tooley et al., 2024). The number of Oregon participants is similar to the one used in other brain map-based analyses of clinical and neurotypical populations (Petrican et al., 2025a,b). The selection criteria for the HCP reference group are detailed in section 2.3 and in the SI.

The generalizability of all the CCA and PLS models was evaluated via a 10-fold cross-validation procedure. Permutation and bootstrap sampling (100,000 samples) were used to estimate the statistical significance and reliability of the multivariate model outputs. The robustness of the CCA and PLS models was further probed a number of technical replications, as detailed in section 2.4 and the SI.

## 3. Results

### 3.1 Neonatal alignment with sex-typical brain architecture predicts parental ratings of psychological ill-health in toddlerhood

We started by investigating whether neonatal alignment with sex-typical FC patterns could foreshadow affective and/or behavioural problems in toddlerhood. To this end, we conducted a PLS analysis between regionally specific indices of alignment with female vs male FC and scores on the 5 DSM-oriented CBCL subscales at 18 months. The brain-behavior LV pair identified in the discovery PLS analysis (*p* =.0005, 32% shared variance) was successfully cross-validated across the 10 test folds (*r*_CV_ =.15, permutation-based *p* =.004, 95% confidence interval [CI] = [.05;.24]). The cross-validation tests controlled for parental race, maternal age and education level, the participant’s sex assigned at birth, fetal age at the neonatal scan (in days) and age at the 18-month follow-up (in weeks), congenital abnormalities (De Asis-Cruz et al., 2018; Provost et al., 2023) (N = 10 neonates), admittance at the neonatal intensive care unit (NICU) (N = 95 neonates), sedation (Greicius et al., 2008; Martuzzi et al., 2010) (N = 3 neonates) and number of motion-related noise components extracted during preprocessing as a participant-level motion metric (see SI).

Next, we examined the correlations (i.e., cross-loadings) between the cross-validated brain LV and each of the 5 DSM-oriented CBCL scales after controlling for the overlap among the latter (*r*s from.37 to.51). This revealed that the identified brain-behavior LV pair reflects the relationship between anxiety at 18 months and neonatal alignment with female-typical FC patterns (cross-loading_CV_ =.12, 95% confidence interval [CI] = [.02;.22]), as well as the link between ASD-relevant challenges in toddlerhood and neonatal alignment with male-typical FC patterns (cross-loading_CV_ =.11, 95% confidence interval [CI] = [.01;.19]) (Figure 2-A, D). Both patterns of alignment with sex-typical FC emerged primarily in default mode (DMN) and Control, and, to a lesser extent, Dorsal Attention (DAN) and Salience/Ventral Attention (SAL/VAN) regions (Figure 2-B,C, E,F).

**Figure 2.**
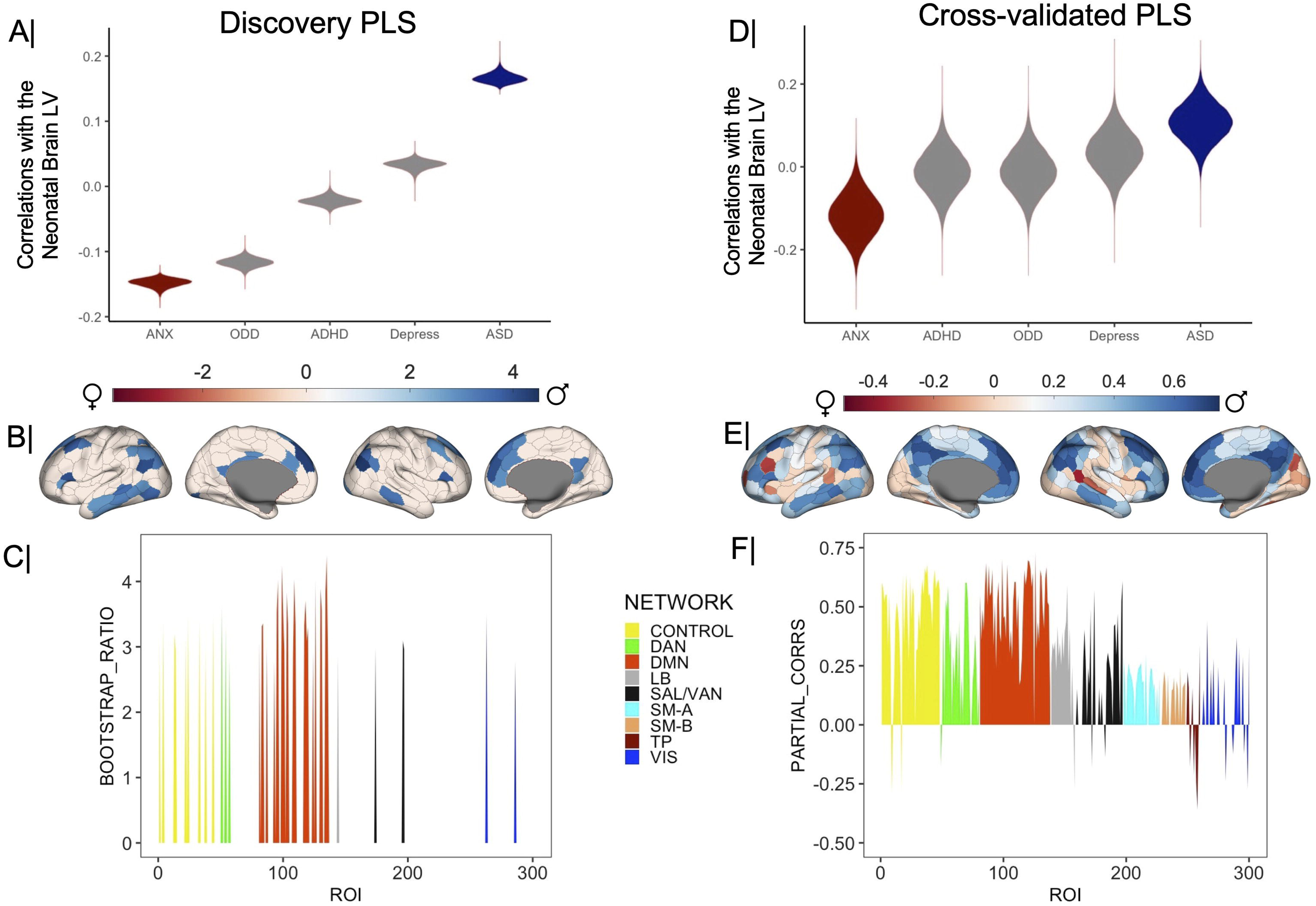
Neonatal alignment with sex-typical FC and psychiatric risk in toddlerhood. The correlation of the neonatal brain LV scores with psychiatric disorder scores in the discovery (panel A) and cross-validated (panel D) PLS analysis. These are partial correlation coefficients controlling for the overlap among the psychiatric disorder scores and the confounders listed in the corresponding Results section. The violin plots in panels A and D depict the distribution of these correlation coefficients across the 100,000 bootstrap samples from the discovery or cross-validated PLS analysis. In these panels, gray colored violin plots indicate a non-robust correlation between the psychiatric disorder scores and the discovery or predicted brain LV scores across all participants. Panel B depicts the parcel-specific weights/loadings on the brain LV identified with the discovery PLS analysis with a bootstrap ratio greater than 2.75 in absolute value (equivalent to a 99% CI). Panel C presents the same information aggregated by the Schaefer atlas networks. Panel E depicts the Schaefer parcels robustly correlated (based on cross-validated 99% CIs) with the predicted value of the brain LV from the cross-validation procedure. These are partial correlations controlling for the confounders listed in the corresponding Results section. Panel F presents Schaefer network-based distributions of partial correlations summarizing the parcel-specific results from panel (E). The subnetworks from the Schaefer 17-network atlas (e.g., Control A/B/C) have been combined into one to increase comparability with the Gordon atlas. In panels C and F, positive values indicate greater alignment with male-, rather than female-typical FC, whereas negative values indicate greater alignment with female-, rather than male-typical FC for a specific parcel. As stated in the main text, alignment with sex-typical FC patterns was estimated in reference to resting state connectivity data from the Human Connectome Project. BSR = bootstrap ratio. FC = functional connectivity. PLS = partial least squares. LV= latent variable. CI = confidence interval. Schaefer networks: TP= temporo-parietal. SAL-VAN = salience/ventral attention. LB = limbic. DMN = default mode. DAN = dorsal attention. SM-A = somatomotor-A. SM-B =somatomotor-B. VIS = visual.

### 3.2 Epigenetic age, cognitive enrichment and maternal mood modulate the predictive power of neonatal FC for autism spectrum problems at 18 months

Analysis 1 established that complementary patterns of neonatal alignment with sex-typical brain architecture relate to anxiety vs ASD-relevant problems in toddlerhood. Next, we examined maturational and environmental factors that could exacerbate or weaken these effects. To this end, we conducted two CCAs linking anxiety (Analysis 2a, Figure 1-J) or ASD-relevant problems (Analysis 2b, Figure 1-K) to the cross-validated brain LV from Analysis 1, indices of neonatal maturation (epigenetic age, gestational age at birth [in days]), maternal medical problems and features of the home environment (maternal depression and anxiety, cognitive enrichment) reported by parents at the 18-month follow-up. In the cross-validation of the CCAs, we controlled for scores on the remaining DSM-oriented CBCL subscales, detection success rate for the DNA methylation signal, parental race, maternal age and education level, the participant’s sex assigned at birth, fetal age at the neonatal scan (in days) and at the 18-month follow-up (in weeks), congenital abnormalities, NICU admittance, sedation and number of motion-related noise components extracted during the fMRI preprocessing.

Unlike Analysis 2-a (Figure 1-J, *r*_CV_ =.04, permutation-based *p* =.452), Analysis 2-b yielded a CCA mode that was successfully cross-validated (Figure 1-K, *r*_CV_ =.15, permutation-based *p* =.003, 95% confidence interval [CI] = [.03;.25], Figure 3-B). The results indicated that greater neonatal alignment with male-typical FC patterns, a younger epigenetic age and poorer maternal health (i.e., higher depression and anxiety, more numerous medical problems) exacerbate, whereas higher cognitive enrichment at home alleviates risk for ASD-relevant problems in toddlerhood (Figure 3-A). The near-zero correlations of neonatal brain LV and epigenetic age with one another (*r* of-.01) and with the remaining variables (*r*s from-.05 to.05) imply an additive effect of the neurobiological and environmental variables to risk for ASD-related difficulties in toddlerhood. Moreover, the modest association of home enrichment with maternal mood (*r*s of-.10 [anxiety] and-.20 [depression]) and maternal medical history (*r* of.02) suggest that the beneficial effect of cognitive stimulation is distinguishable from that of maternal mental and physical health.

**Figure 3.**
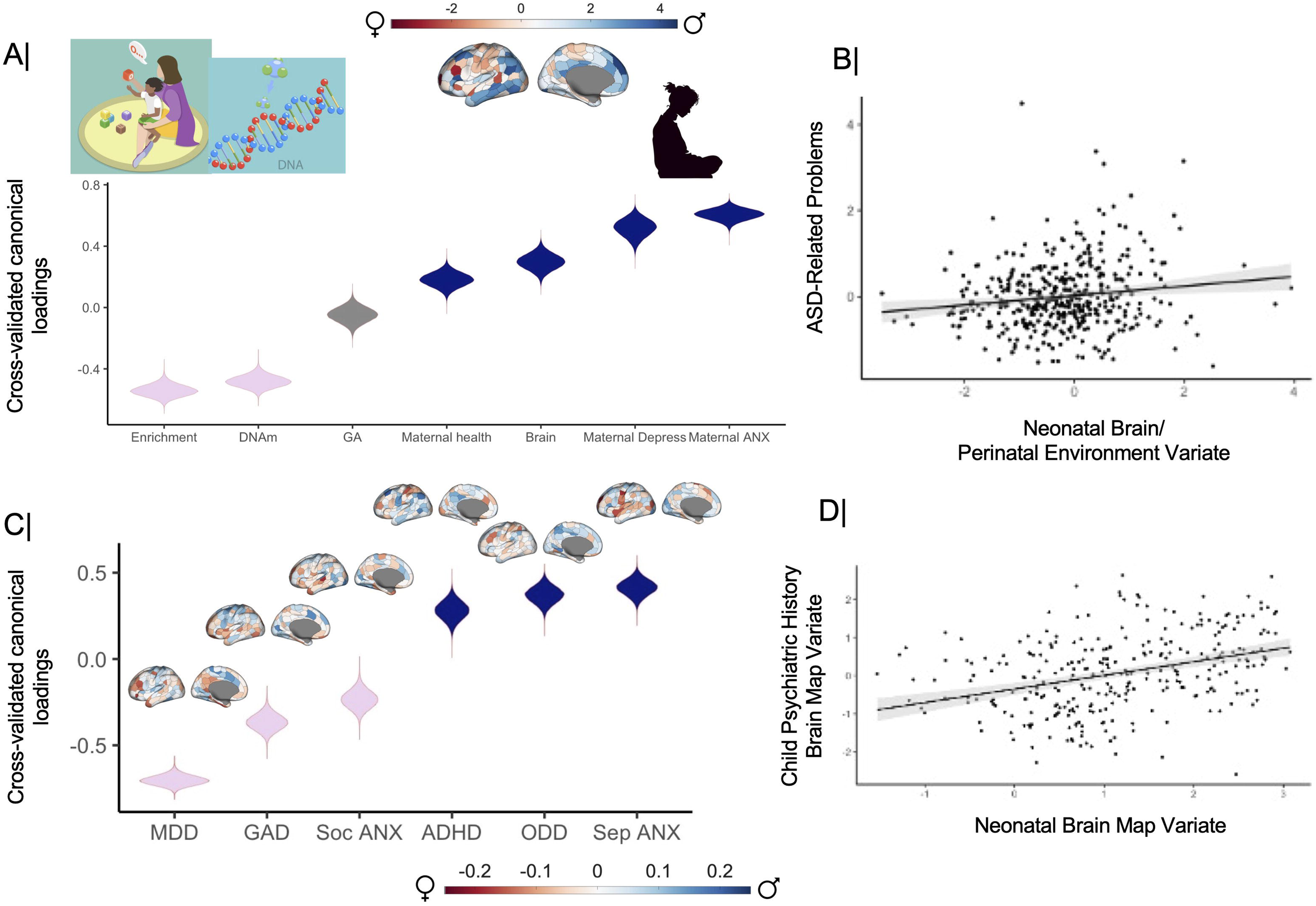
Alignment with Sex-Typical FC in Toddlerhood and Childhood: Links with Environmental Variables and Mental Health. The partial correlation coefficients describing the relationship of the cross-validated neonatal brain LV from Analysis 1, measured infant (DNAm, GA), maternal (physical health, depression, anxiety) and home environment (cognitive enrichment) variables with the predicted value of their corresponding canonical variate across all test CCA folds are presented in panel A (dHCP sample). Panel B contains the scatter plot describing the linear relationship between ASD-relevant symptoms in toddlerhood (y-axis) and the CCA variate pictured in panel A, which captures the additive effects of neonatal alignment with male-typical FC, neonatal epigenetic age and features of the perinatal environment (x-axis) (after controlling for the overlap with the other DSM-oriented CBCL scales and the confounders listed under “Control variables for the dHCP and Oregon samples”). Panel C shows the correlation coefficients describing the relationship between the brain maps quantifying alignment with sex-typical FC in connection to psychiatric history and the predicted value of their corresponding canonical variate across all test CCA folds (Oregon sample). Patterns of alignment with sex-typical FC corresponding to each disorder are pictured above the corresponding violin plot. Panel D contains the scatter plot describing the linear relationship between the cross-validated and unthresholded neonatal brain LV map from Analysis 1 and the CCA variate pictured in panel C. The violin plots in panels A and C depict the distribution of (partial [A only]) correlation coefficients across the 100,000 bootstrap samples from the cross-validation procedure. Gray colored violin plots indicate non-robust canonical loadings (based on the bootstrapped 99% CIs from the cross-validation procedure). The shaded area in panels B and D constitute the 95% CI for the correlation coefficient. CCA = canonical correlation analysis. CI = confidence interval. DNAm = DNA methylation (based on which epigenetic age was estimated, see text for details). GA = gestational age. ANX = anxiety. ADHD = attention deficit hyperactivity disorder. ASD = autism spectrum disorder. MDD = major depressive disorder. Sep = separation. GAD = generalized anxiety disorder. ODD = oppositional defiant disorder. Soc = social.

### 3.3 Neonatal alignment with sex-typical FC along the unimodal-to-transmodal axis and in areas of greater cholinergic receptor density

Next, we sought to situate the identified neonatal profile in the context of canonical multilevel models of brain maturation. Hence, we conducted a CCA relating the cross-validated brain LV from Analysis 1 (Figure 2-E) to the first two functional gradients (gradient 1: unimodal-to-transmodal, gradient 2: visual-to-sensorimotor) identified by Margulies et al. (2016), as well as the receptor and transporter density maps for the major neurotransmitter systems (i.e., serotonin (5-HT), dopamine [DA], acetylcholine [Ach], GABA, glutamate [GLU] and norepinephrine [NE]) described by Hansen et al. (2022). This analysis yielded a sole CCA mode which was successfully cross-validated (*r*_CV_ =.69, *p_spin_* = 10^-5^, 95% confidence interval [CI] = [.61;.74]) and indicated that ASD-relevant alignment with male-typical FC and anxiety-linked alignment with female-typical FC emerge primarily in transmodal (rather than unimodal) areas, particularly those higher in A4 cholinergic receptor density and lower in 5-HT, DA, Ach, and NE transporter density (Figure 4).

**Figure 4.**
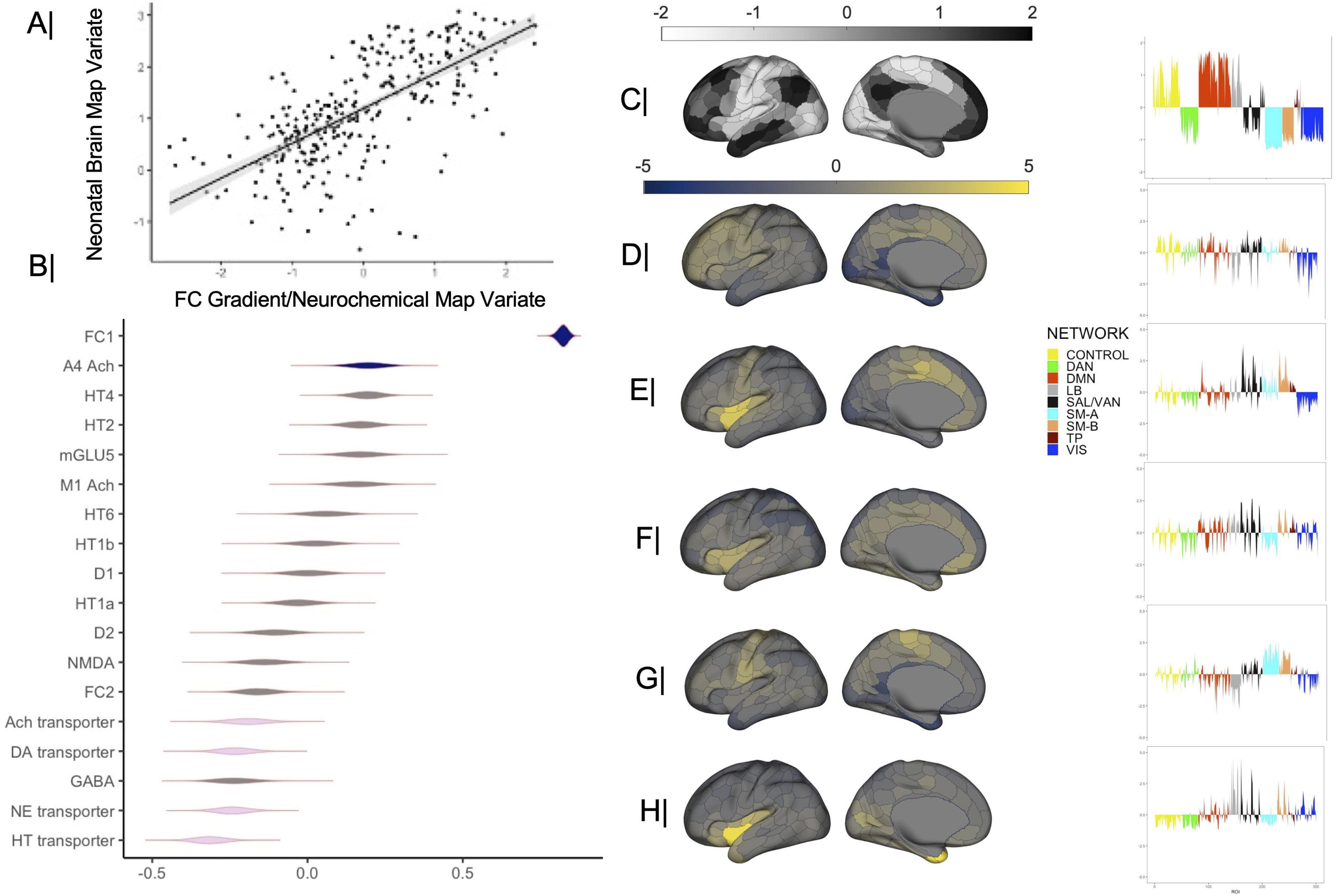
Neonatal alignment with sex-typical FC: Correlation with functional hierarchies and neurochemical architecture. Panel A contains the scatter plot describing the linear relationship between the cross-validated neonatal brain LV (Figure 2-E) and the CCA variate pictured in panel B. The shaded area in panel A constitutes the 95% CI for the correlation coefficient. Panel B shows the correlation coefficients describing the relationship of the functional gradient and receptor/transporter density maps with the predicted value of their corresponding canonical variate across all test CCA folds. The violin plots depict the distribution of correlation coefficients across the 100,000 bootstrap samples from the cross-validation procedure. Gray colored violin plots indicate non-robust canonical loadings (based on the bootstrapped 99% CIs from the cross-validation procedure). Panels C-H depict the relative distribution (z-scored across all the parcels in the Schaefer atlas) at the level of each parcel (left side) or Schaefer network (right side) for the somatomotor-to-transmodal functional gradient (C), A4 Ach receptor (D), Ach transporter (E), DA transporter (F), NE transporter (G) and HT transporter (H). LV = latent variable. CI = confidence interval. (5-)HT =serotonin. Ach = acetylcholine. D= dopamine. GABA = gamma-aminobutyric acid; GLU = glutamate. NMDA = N-methyl-D-aspartate receptor. NE = norepinephrine. Schaefer networks: TP= temporo-parietal. SAL-VAN = salience/ventral attention. LB = limbic. DMN = default mode. DAN = dorsal attention. SM-A = somatomotor-A. SM-B =somatomotor-B. VIS = visual.

### 3.4 Robust overlap between the neonatal and child profiles of alignment with sex-typical FC linked to mood, ADHD and behavioural problems

Lastly, we tested whether the neonatal profile of alignment with sex-typical FC linked to anxiety and ASD-relevant problems in the dHCP sample would predict childhood psychiatric history in the Oregon ADHD 1000 cohort (Nigg et al., 2023). A CCA linked the cross-validated neonatal brain LV from Analysis 1 (Figure 2-E) to the 6 brain maps quantifying alignment with sex-typical FC in connection to a child history of MDD, anxiety (separation, social, GAD), ODD and current diagnosis of ADHD. This analysis yielded a sole CCA mode which was successfully cross-validated (*r*_CV_ =.36, *p_spin_* = 2 x 10^-5^, 95% confidence interval [CI] = [.26;.45], Figure 3-D). The results indicated a robust spatial overlap between neonates and children in corresponding patterns of alignment with sex-typical FC. Thus, on one hand, alignment with female-typical FC correlated with anxiety (social, GAD) in both samples, and, in childhood only, also a lifetime history of MDD.

Complementarily, alignment with male-typical FC related to ASD-relevant problems in toddlerhood, while in childhood it was connected to currently diagnosed ADHD and lifetime history of ODD and separation anxiety (Figure 3-C).

## 4. Discussion

The past two decades have seen a growing interest in shifting from a reactive to a pre-emptive approach to psychiatric disorders (Insel, 2007) by integrating insights from (epi)genetics, neuroscience and cognitive science (Cuthbert & Insel, 2013). To answer these global calls for a redefinition of mental health in a multi-level framework spanning cells to functional systems (Insel et al., 2010), our present investigation combined neuroimaging, epigenetic, clinical and environmental assessments from a neonatal (N = 411) and a child sample (N = 214).

Consistent with the well-documented role of association networks in neurodevelopment (Li et al., 2023; Sydnor et al., 2021; Yin et al., 2025) and sex-based classification (Keller et al., 2025; Serio et al., 2024; Shanmugan et al., 2022), our analyses unveiled complementary patterns of alignment with sex-typical FC which unfolded along the S-A axis and predicted distinguishable psychiatric profiles, broadly replicated between toddlerhood and late childhood. Thus, across sexes, greater neonatal alignment with female-typical FC patterns, particularly for DMN and Control regions, forecasted higher anxiety at 18 months, whereas the same brain pattern correlated with lifetime history of MDD and anxiety (social, GAD) among the Oregon youths. These results extend recent reports in which stronger adolescent alignment with female-typical FC in association areas related to greater internalizing symptoms across sexes (Petrican et al., 2025a). Here, we show that these relationships emerge within the first year of life. Our present findings and those reported by Petrican et al. (2025a) also indicate that relative alignment with female-typical FC along the S-A axis could aid cross-sectional detection of both anxiety and depression, but longitudinal prediction of anxiety only.

Conversely, across sexes, greater neonatal alignment with male-typical FC patterns, especially in DMN and Control regions, foreshadowed more severe ASD-relevant problems in toddlerhood, whereas in late childhood, it correlated with a current diagnosis of ADHD and lifetime history of ODD and separation anxiety. The link between ASD and alignment with male-typical FC along the S-A axis resonates with recent proposals that alterations in the S-A functional hierarchy could provide a systems-level explanation for the heterogeneity of challenges associated with ASD (Bernhardt et al., 2025; Hong et al., 2019). Taken together, our findings imply that early life hyper-alignment with the adult S-A axis profile typical of one’s own sex could underpin vulnerability to ASD in boys and anxiety in girls, whereas divergence may heighten vulnerability to ASD-linked challenges in girls and anxiety in boys.

The relationship between ASD and neonatal alignment with male-typical FC patterns dovetails with the reported greater prevalence of ASD among males (Lord et al., 2020) and observations that some autistic traits could reflect “male-typical” behaviors (Baron-Cohen, 2009). The ASD-predictive brain pattern also intersected the male-typical FC maps tracking current ADHD diagnosis and lifetime diagnoses of ODD and separation anxiety in childhood. These findings imply that alignment with male-typical FC along the S-A axis could contribute to the pathophysiological resemblance between ASD and ADHD (Itahashi et al., 2025; Mikami et al., 2019), as well as the clinical overlap between these two conditions and ODD with regards to social-affective processes (ASD-ODD(Sun et al., 2022; Zhang et al., 2024)) and attention (ADHD-ODD(Baskin-Sommers & Brazil, 2022; Baskin-Sommers et al., 2022)). Our observed neural overlap between ASD-relevant problems in toddlerhood and separation anxiety in childhood fits with existing evidence on the relevance of separation anxiety (but not other types of anxiety) to understanding insistence on sameness in ASD (Black et al., 2017; Rodgers et al., 2012).

In the dHCP sample, there was suggestive evidence that characteristics of the infant, the mother and of the home environment can independently influence the predictive power of neonatal brain architecture for psychiatric symptoms in toddlerhood. Specifically, neonatal alignment with male-typical FC patterns was most predictive of ASD-relevant problems among those toddlers who were epigenetically less mature at birth, had less cognitively stimulating home environments and mothers who reported poorer physical and mental health. The detrimental contribution of younger epigenetic age is compatible with existing theories linking ASD to prolonged neurocognitive development (Johnson et al., 2015), whereas the alleviating effect of home enrichment fits with cross-species findings that cognitively stimulating environments can protect against the sequelae of early life adversity exposure (Wang et al., 2025; You et al., 2025). The differential relevance of home enrichment to ASD, but not anxiety underscores the value of an “embedded brain” approach in personalizing mental health interventions through consideration of the reciprocal effects between intra-individual (e.g., neurocognitive traits) and environmental (e.g., family/peer relationships, cognitively stimulating activities) factors (Viding et al., 2024). Our results further imply that the effectiveness of parenting and teacher-related interventions targeting ASD (Gora et al., 2024) and related conditions, such as ADHD (Shephard et al., 2022) and ODD (Sethi et al., 2023), could be enhanced by combining social interactions with mentally stimulating activities, as implemented in dialogic book-sharing (Boyle et al., 2019; Murray et al., 2016; Murray et al., 2023).

The complementary patterns of alignment with sex-typical FC, linked to anxiety vs ASD-relevant problems in toddlerhood, robustly overlapped areas richer in nicotinic cholinergic receptors. Cholinergic signalling enhances the precision of bottom-up signals in cortical hierarchies (Shine et al., 2021) and contributes to attention and encoding of emotionally salient information (Ananth et al., 2023), as well as cross-species responses to environmental adversity (Mineur et al., 2022), including exposure to prenatal maternal stress (Vaknine Treidel et al., 2025). In light of this literature, our findings raise the possibility that interactions between the cholinergic system and sex hormone metabolic processes (Newhouse & Dumas, 2015) may give rise to the complementary brain patterns observed for ASD vs anxiety. Specifically, greater alignment with male-typical FC patterns, particularly in DMN and Control regions, may be relevant to understanding the putative devaluation of prior knowledge and overweighting of sensory input in ASD (Bernhardt et al., 2025), whereas the alignment of these regions with female-typical FC patterns could help elucidate the neurobiological underpinnings of the cross-domain divergence in cognition and social processing observed in anxiety disorders (Akiki et al., 2025).

The present research has several limitations which could though act as a springboard for future work in this area. First, although we provided evidence that home enrichment and maternal health can influence the predictive power of neonatal FC for ASD-relevant problems in toddlerhood, our analyses could not elucidate underlying mechanisms. Second, we did not investigate the potential moderating role of pubertal timing in the Oregon sample. However, this line of inquiry is worth pursuing given the well-documented contribution of pubertal timing to transdiagnostic psychiatric risk (Hamlat et al., 2019) and the relevance of pubertal hormones to neural sexual differentiation (Piekarski et al., 2023; Schulz & Sisk, 2016). Third, we used two independent samples to estimate the stability of the relationship between alignment with sex-typical FC patterns and mental health symptoms from birth to late childhood. The underlying assumption, supported by the data (Figures S5-6, A-F), was that alignment with sex-typical FC profiles emerges gradually from birth onwards, with young adults constituting the most appropriate reference group because they show maximal sexual differentiation. Accordingly, while female-to-male FC strength gradients based on the dHCP or Oregon data as a reference performed well within-group (Figures S5-6, G, H), they could not be validated across age groups (permutation-based *p*-values >.27). Longitudinal samples are though needed to characterize normative trajectories of neural sexual differentiation and to establish whether the same regional patterns of alignment with sex-typical FC are persistently linked to broadly similar psychiatric symptoms from toddlerhood onwards.

In sum, we identified complementary patterns of neonatal alignment with sex-typical brain architecture which overlapped regions rich in nicotinic cholinergic receptors and, at 18 months, distinguished between mental health problems with sex-biased prevalence. We also provided evidence that the predictive power of alignment with male-typical FC for ASD-linked behaviors is impacted by the infant’s epigenetic age around birth, maternal health and cognitive enrichment at home. We further demonstrated that, among children, patterns of alignment with sex-typical FC, similar to those observed in neonates, differentiate a history of MDD and anxiety (social, GAD) from a current diagnosis of ADHD, as well as a history of ODD and separation anxiety. By linking regional alignment with female-vs male-typical FC to similar psychiatric outcomes across sexes in toddlerhood and childhood, and documenting their susceptibility to adjustable environmental conditions, our research seeks to act as a springboard for future precision medicine efforts to optimize early life maturation and psychological well-being.

## Materials & Correspondence

Correspondence and material requests should be addressed to R.P.

## Data statement

The raw data are available at https://nda.nih.gov/study.html?id=1938 (DOI: 10.15154/1528485) (Oregon), at https://db.humanconnectome.org (HCP) and at https://nda.nih.gov/edit_collection.html?id=3955 (dHCP) upon completion of the relevant data use agreements. The dHCP data used in this report came from Data Release 3.0 (DOI: 10.15154/92vw-g837).

## Code availability

We used already existing code, as specified in the main text with links for free download.

## Acknowledgments

Data used in the preparation of this article were obtained from the Oregon ADHD-1000: A longitudinal data resource enriched for clinical cases and multiple levels of analysis, Human Connectome Project, WU-Minn Consortium (Principal Investigators: David Van Essen and Kamil Ugurbil; 1U54MH091657; funders: the 16 NIH Institutes and Centers that support the NIH Blueprint for Neuroscience Research and the McDonnell Center for Systems Neuroscience at Washington University), as well as from the Developing Human Connectome Project (dHCP). The Oregon ADHD-1000 project was supported by multiple NIMH grants: 1R01-MH59105 (Nigg); R01-MH63146 (Nigg); R01-MH070004 (Friderici); 1R01MH86654 (Nigg-Fair); R01MH099064 (Nigg); 1R37MH59105 (Nigg); 2R56MH086654 (Nigg); R01MH115357 (Fair-Nigg); 2R37MH59105 (Nigg); R01 MH096773 (Fair); K23 MH108656 (Karalunas). Data storage at OHSU is supported by the Oregon Clinical and Translational Research Institute funded by a grant from the National Center for Advancing Translational Sciences (NCATS), National Institutes of Health, through Grant Award Number UL1TR002369. The Developing Human Connectome Project was funded by the European Research Council under the European Union Seventh Framework Programme (FP/20072013)/ERC Grant Agreement no. 319456 and was also supported by the NIHR Biomedical Research Centres at Guys and St Thomas NHS Trust. This manuscript reflects the views of the authors and may not reflect the opinions or views of the NIH, HCP, dHCP or Oregon ADHD 1000 consortium investigators.

